# Environmental sex determination in the cyst nematode *Globodera pallida* defaults to male development

**DOI:** 10.1101/2025.11.04.686318

**Authors:** Arno S. Schaveling, Stefan J.S. van de Ruitenbeek, Geert Smant, Mark G. Sterken

## Abstract

Environmental sex determination (ESD) enables organisms to adjust their sexual fate in response to external cues. Fluctuating sex ratios have long suggested the presence of ESD in populations of plant-parasitic nematodes. We show that in the potato cyst nematode *Globodera pallida* sex is regulated by nutritional cues. By manipulating sucrose availability to the host plant, we could steer the sex determination of *G. pallida*. Whereas high-sucrose medium promotes female development, low-sucrose medium promotes male development. Transcriptome analyses on the early stages of parasitism reveal that female development requires extensive transcriptional activation and post-transcriptional regulation. We identify *Gp-lin-29*, a transcription factor homologous to *lin-29* in *Caenorhabditis elegans*, as a potential regulator of ESD. Small RNA sequencing uncovered the male-biased expression of *Gp-let-7*, a putative repressor of *Gp-lin-29*, and the female-biased expression of four miRNAs, including *Gp-miR-100*, located at the same genomic locus as *Gp-let-7*. Target prediction and enrichment analyses suggest that these female-biased miRNAs actively suppress male developmental programs. Together, our findings support a model in which *G. pallida* juveniles follow a default male developmental trajectory unless redirected by favourable environmental cues to become female. This study provides mechanistic insight into ESD in cyst nematodes and positions *G. pallida* as a tractable system for exploring epigenetic regulation of developmental plasticity.

## INTRODUCTION

Sex determination mechanisms vary widely within the animal kingdom. These mechanisms are typically categorized as either genetic sex determination (GSD) or environmental sex determination (ESD), depending on the cue that triggers sex differentiation (Uller et al., 2007). In the case of GSD, sex is determined by a gene or a set of genes generally located on sex chromosomes. In ESD, environmental factors such as temperature and nutrient availability determine sex. While mammals appear to be restricted by GSD, other animals such as fish, reptiles, and amphibians can adjust their sex based on environmental and social cues (Navara, 2018). Interestingly, both GSD and ESD are present within the phylum Nematoda. For instance, the model species *Caenorhabditis elegans* exemplifies GSD, where the number of X chromosomes determines the sex, resulting in XX hermaphrodites and XO males (Zarkower, 2006). In contrast to GSD, the sex determination of certain nematode species parasitizing on arthropods, vertebrates, and plants have shown to be controlled by infection densities and nutrient availability (Haag, 2005). Despite numerous observations of ESD in nematodes, the underlying mechanism remains elusive.

Cyst nematodes are highly adapted and specialized obligate plant-parasites (Goverse & Mitchum, 2022). Members of the cyst nematode genera *Heterodera* and *Globodera* cause major yield losses in crops, including cereals, vegetables, sugar beets, and potatoes. It has been known for over a century that environmental factors influence the sex ratios of cyst nematodes (Müller & Molz, 1920; Trudgill, 1967). Particularly, nutrient availability plays a crucial role in sex ratios of *H. schachtii*, *G. pallida,* and *G. rostochiensis*. This can be observed indirectly by variation in infection densities, sucrose levels, and syncytium quality (Anjam et al., 2020; Grundler et al., 1991; Hofmann & Grundler, 2006; Trudgill, 1967). Also plant resistance responses are known to increase the male-to-female ratios of cyst nematodes (Anwer et al., 2018; Caromel et al., 2005; Castelli et al., 2005; Fournet et al., 2018; Rice et al., 1985; Schouten, 1993; Sobczak et al., 2005; Van Der Vossen et al., 2000). As the production of hundreds of eggs is highly nutrient-demanding (Muller et al., 1981), limited nutrient availability might trigger a mechanism preventing juveniles from developing into females.

Epigenetic regulation is a candidate mechanism driving ESD. Core epigenetic factors include histone modifications, DNA methylation, and regulatory non-coding RNAs. Non-coding RNAs have shown to play a key role in the adaptation to environmental conditions in free-living and cyst nematodes (Fusca et al., 2022; Ste-Croix et al., 2023). Moreover, small non-coding RNAs (sRNAs) have shown crucial regulatory functions in sex determination of silkworms (Kiuchi et al., 2014), Oriental fruit flies (Peng et al., 2020), and Reeves’ pond turtles (Xiong et al., 2020). In *C. elegans*, sRNAs tightly regulate the somatic sex determination pathway (McJunkin & Ambros, 2017). An example of which is the sRNA-mediated regulation of the transcription factor *tra-1*, the master regulator of X-chromosome dosage compensation and sex determination (Tang et al., 2018). Given the challenges of studying ESD in larger organisms, cyst nematodes could serve as a valuable model system for investigating epigenetic mechanisms involved in ESD.

In our study, we established experimental conditions to steer the environmentally-regulated sex determination of the potato cyst nematode *G. pallida*. Juveniles inoculated on potato cuttings cultured on low-sucrose medium showed a 41-fold reduction in the number of adult females and a 3-fold increase in the number of adult males compared to juveniles inoculated on potato cuttings grown on high-sucrose medium. We used these experimental conditions to investigate the mechanism underlying ESD. Transcriptome sequencing of the early stages of parasitism revealed extensive transcriptional activation and post-transcriptional regulations during female development, including the female-biased expression of a *lin-29*-like transcription factor. Small RNA sequencing revealed the male-biased expression of *let-7*, known to suppress *lin-29* in *C. elegans*, and the female-biased expression of four miRNAs. MiRNA target prediction of these four miRNAs suggests that these miRNAs target genes involved in male-development. Finally, we argue why the default developmental trajectory leads towards males.

## METHODS

### Manipulating sex determination by differences in host sucrose availability

*In vitro* infection assays were performed as described by Goverse et al. (2000) with additions made by Zheng et al. (2022) and (Schaveling et al., 2025). In brief, cuttings of the susceptible potato variety Desirée were grown on Gamborg B5 medium supplemented with specified amounts of sucrose. Centimetre long one-node cuttings were grown with one per square (12×12cm) plate. After 14 days, cuttings were inoculated with 100 surface-sterilized *G. pallida* E400 Rookmaker pre-parasitic second-stage juveniles (ppJ2s) per plate. Male and female development were tracked over time with an Olympus SZX10 binocular microscope (Olympus, Japan) with a × 1.5 objective and ×3.2 magnification. Pictures were taken with an AxioCam 712 colour camera (Zeiss, Germany). For visual purposes, contrast and clarity of all photos were enhanced in Adobe Lightroom (v8.1; http://adobe.com/products/photoshop-lightroom.html). Supplementary video 1 was created in PowerPoint for Microsoft 365 (v16.0.1).

The mRNA and small RNA sequencing experiment was conducted in four time-separated batches. Each batch started with a total of 120 to 130 plates. Half of the plates contained 20 g/L sucrose, the other half contained 1.5 g/L of sucrose. Upon inoculation, a subsample of the ppJ2s was taken for RNA extraction. Infected root segments were harvested at 1, 3, 6, and 9 days post inoculation (dpi). At 1 and 3 dpi, infected root segments of 15 plants were pooled per sample and at 6 and 9 dpi, infected root segments of 10 plants were pooled per sample. All uncontaminated plates that were not used for RNA extraction were used for scoring of males and females at 35 dpi. A total of 65 plates was scored across four batches. Total RNA was extracted with the Maxwell 16 LEV-plant RNA kit (Promega, USA), according to the manufacturer’s instructions. Samples were split, to send ≥300 ng of total RNA for transcriptome sequencing and ≥1ug of total RNA for sRNA sequencing to BGI Genomics (Hong Kong, China). For transcriptome sequencing, stranded libraries were sequenced on the DNBseq G400 platform, generating over 50 million paired-end 150 base pair reads per sample (**Supplementary table 1**). Two samples of batch 4 (B_4.2_1,5; B_4.3_1,5) failed at quality control and were not taken along for transcriptome sequencing. Small RNA samples were pre-treated with RNA 5’ Polyphosphatase (Biosearch™ Technologies, UK). For sRNA sequencing stranded libraries were also sequenced on the DNBseq platform, generating over 20 million single-end 50 base pair reads per sample (**Supplementary table 2**).

### General notes on data analysis

All custom scripts and data are available through gitlab (https://git.wur.nl/published_papers/Schaveling_2025_ESDpallida). Unless described otherwise, data was analyses with R (v4.3.2; R Core Team, 2013) and R studio (v2023.06.1; Team, 2015) using the *ggplot2* package (Villanueva & Chen, 2019) for data visualisation. Other packages are mentioned at the relevant sections. Differential expression analyses were performed with DESeq2 (Love et al., 2014) via the SeqMonk interface (v.1.48.1, https://www.bioin formatics.babraham.ac.uk/projects/seqmonk/). Small RNA sequencing data is deposited at BioStudies (E-MTAB-15069). Structural annotation of sRNA loci is deposited at the *G. pallida* Rookmaker genome at the NCBI (PRJEB91928).

### Processing transcriptome sequencing data

Trimmed reads were obtained from BGI (Hongkong). This yielded an average of 104 million clean reads per sample, with a Q30 ranging from 91.7% to 94.9% (**Supplementary table 1**). Samples were analysed using MultiQC (Ewels et al., 2016). The reads were mapped against the *G. pallida* Rookmaker genome (v.1.0; Schaveling et al., 2025) using hisat2 (v.2.2.1; Kim et al., 2019). For the ppJ2 samples, between 87.0% and 88.7% of reads mapped and correctly paired to the Rookmaker genome. On average a sample showed expression for 89% of the genes. Among the infected root tissue samples we identified two samples that failed quality control. Hierarchical clustering of pairwise Pearson correlations based on the log_2_(TPM) revealed that two samples (both samples of batch 4 at 1 dpi) clustered poorly with the others (**Supplementary Figure 2**). So these two samples were excluded from expression analysis. For the root tissue samples that passed quality control the reads correctly mapping and pairing varied between 0.9% and 24.7%, on average detecting expression for 72% of all genes. BAM files were loaded into SeqMonk. Raw count values and transcript per kilobase million (TPM) values were generated using the RNA-Seq quantitation pipeline in SeqMonk and extracted for further analysed in R.

### Transcriptome analyses identify sex-biased expression patterns

For gene-centric analyses, the ratio between the TPM and the mean TPM (TPM_ratio_) was calculated for each transcript by:

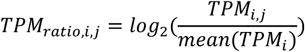

where *TPM_i,j_* was the TPM value of the gene *i* (one out of 21,026 genes from *G. pallida*) and sample *j* (one out of 36 samples) and *mean(TPM_j_)* was the mean TPM within gene *i*. Principal component (PC) analysis was performed on the TPM_ratios_, using the R function prcomp.

To identify differentially expressed genes (DEGs) with DESeq2, raw transcript count values were compared between male and female development at each of the four post-infection timepoints. We applied independent filtering and correcting p-values for multiple testing by the Benjamini-Hochberg method (false discovery rate; FDR). The UpSet plot was made with the *ComplexUpset* package (Michał Krassowski et al., 2022).

K-means clustering was used to identify similar expression patterns among sex-biased genes. Based on the TPM_ratios_, we tested clustering with up to 20 clusters using the kclust function from the *broom* package with parameters iter.max = 3,000 and nstart = 1,000.

### Characterization of sex-biased genes

We used OmicsBox (v3.0; Götz et al., 2008) to annotate protein sequences with GO-terms. GO-term enrichment analyses were performed with hypergeometric tests, using the phyper function in R. Resulting p-values were FDR-corrected for multiple testing.

Protein sequences of all sex-biased genes were blasted against the non-redundant protein sequences on the NCBI webserver (https://blast.ncbi.nlm.nih.gov/Blast.cgi) at default settings. The BLASTp hit with the lowest e-value that was not described as “unnamed protein”, “hypothetical protein”, “uncharacterised protein”, or “protein of unknown function” was considered the most informative BLASTp hit (**Supplementary table 4**). Domain predictions on three sex-biased transcription factors were conducted with InterPro (Paysan-Lafosse et al., 2023). Known physical and regulatory *lin-29* interactions in *C. elegans* were obtained at WormBase (Sternberg et al., 2024) on November 5^th^, 2024 (https://wormbase.org/db/get?name=WBGene00003015;class=Gene).

### Processing sRNAseq data and generating structural annotation of sRNA loci

BGI provided an average of 25.3 million clean small RNA reads with an average Q30 of 96.3% (**Supplementary table 2**). All small RNA sequencing data is deposited at BioStudies (E-MTAB-15069). Small RNA reads were processed using ShortStack (v4.0.3; Johnson et al., 2016). First, trimmed reads of the pre-parasitic samples were merged and mapped against the *G. pallida* Rookmaker genome to generate a structural annotation. The resulting GFF3 file is deposited with the Rookmaker genome at NCBI (PRJEB91928). The structural annotation consists of clusters that must pass three criteria: a minimal coverage of 1 RPM, a minimal read depth of 25, and a minimal distance of 75 bp between two clusters. This yielded 11 miRNA clusters, 43,517 siRNA clusters and 2,366 undetermined clusters. Then, all samples were mapped individually and scored on the 45,894 predetermined loci. On average, 78.4% of the ppJ2 sRNA reads mapped to the nematode genome. However, 3% of the reads map to the potato genome. In infected root tissue samples, on average, 13.7% of the reads mapped to the nematode genome, and 8.7% mapped to both the plant and the nematode genome (**Supplementary table 3**). This indicates that a large fraction (64.5%) of the reads mapping to the nematode genome also map to the potato genome. Based on our transcriptome analyses, we again excluded both samples of batch 4 at 1 dpi from further analyses. BAM files generated by ShortStack were analysed in SeqMonk (v1.48.1; www.bioinformatics.babraham.ac.uk/projects/seqmonk). Read length histogram, raw read counts and reads per kilobase million (RPM) values were exported from SeqMonk and analysed in R.

### Small RNAseq data analyses identify sex-biased miRNAs

For principal component analysis, the RPM_ratio_ was calculated for each of the sRNA loci and for each of the samples in the same way as we calculated the TPM_ratio_. Principal component analysis was performed on the RPM_ratio_, as described for the transcriptome data.

To assess the hypothesis of sex-biased *Gp-let-7* expression, differential expression analysis was performed on raw read count values with DESeq2 as described before. However, as we were interested in a specific small RNA locus, we did not correct for multiple testing. To assess whether there were sRNA loci that were consistently differentially expressed during male and female development, we compared all samples representing male development with all samples representing female development with DESeq2, disregarding timepoints. Resulting p-values were FDR-corrected.

### Characterization of sex-biased miRNAs

RNA 2D structures were generated with the Vienna RNAfold web server (http://rna.tbi.univie.ac.at/cgi-bin/RNAWebSuite/RNAfold.cgi) Package 2.0 (Lorenz et al., 2011), with default settings. For miRNA target prediction, we considered the 500 base pair after the stop codon as the putative 3’ untranslated region (UTR). Therefore, we extracted the 500 base pairs downstream of the stop codon of each transcript on the *G. pallida* Rookmaker genome and used this to predict miRNA targets with RNAhybrid (v2.2; Krüger & Rehmsmeier, 2006). Targets were required to have a maximum free energy of -25 kcal/mol and perfect seed matching (nucleotide 2-7 of the mature miRNA sequence). GO-terms that were present in more than two miRNA targets were assessed for enrichment, as described before.

### Conservation analysis of the genetic sex determination pathway of C. elegans

To assess the conservation of the global sex determination pathway of *Caenorhabditis elegans*, protein sequences were retrieved from wormbase.parasite.org (Howe et al., 2017) and ran in a local BLASTp (v2.12.0+) against the protein databases of *G. pallida* Rookmaker genome (Venables & Ripley, 2013), *G. rostochiensis* line 19 and line 22 (van Steenbrugge et al., 2021), and *H. schachtii* IRS (van Steenbrugge et al., 2023), with a cutoff E-value of 1e-5. BLASTp hits were considered homologs when sequence identity exceeded 25% and query coverage exceeded 50%.

## RESULTS

### The sucrose availability for the host affects the sex determination of *G. pallida*

To investigate sex determination mechanisms in *G. pallida*, we determined experimental conditions under which sex could be manipulated. Since sex ratios of the cyst nematode *H. schachtii* could be manipulated by altering the sucrose available for the host (Grundler et al., 1991), with reduced sucrose levels resulting in increased male-to-female ratios, we tested whether sex determination in *G. pallida* depends on host sucrose availability as well. Therefore, we grew *in vitro* potato cuttings on Gamborg B5 medium supplemented with a range of sucrose concentrations. Two-week-old potato cuttings were inoculated with 100 pre-parasitic juveniles (ppJ2s). Sex was scored at 35 days post inoculation (dpi), although, female juveniles could be distinguished from 9 dpi onwards (**Figure 1A; Supplementary video 1**). Over two pilot experiments, we hardly observed female development at sucrose concentrations below 2.5 g/L (**Supplementary Figure 1A&B**). The highest numbers of males were found at 1.5 g/L sucrose (**Supplementary Figure 1B**). At lower sucrose concentrations, also male development was inhibited. For subsequent sex determination experiments, we therefore supplemented the media with either 1.5 or 20 g/L of sucrose.

**Figure 1.**
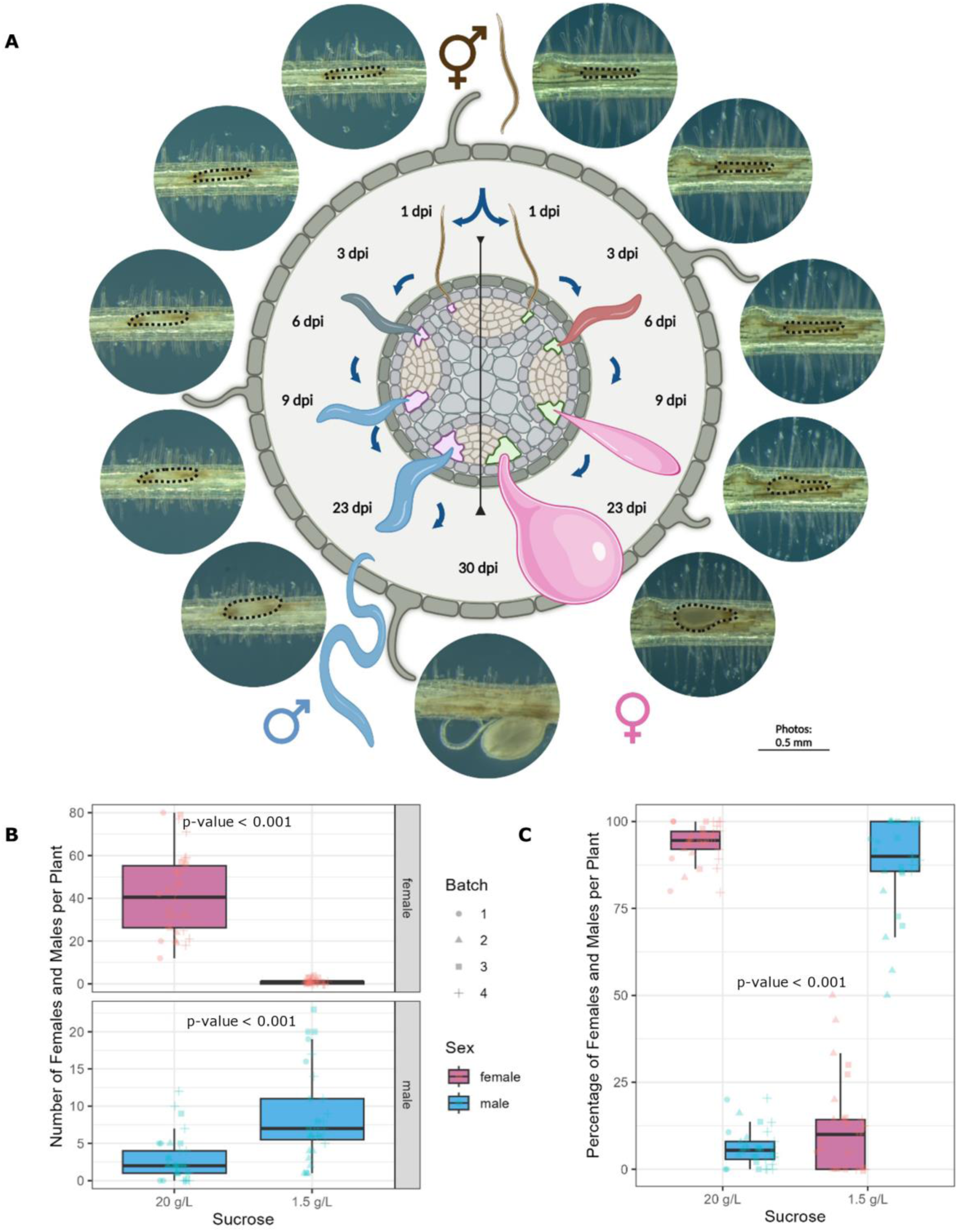
Environmental sex determination in the cyst nematode *Globodera pallida*. **A** Graphical representation of male and female development, surrounded by representative pictures of the corresponding life stages. Second-stage juveniles enter the root (top). Juveniles with limited nutrient availability develop into adult males (left), juveniles with high nutrient availability develop into adult females (right). Dashed lines outline juveniles in the pictures. The figure was made using BioRender (https://BioRender.com/ceroryg). **B** The numbers of males and females per plant, measured over four replicate experiments, show an absolute increase of the number of males on low-sucrose conditions (1.5 g/L sucrose; n_total_ = 62) compared to high-sucrose conditions (20 g/L sucrose; n_total_ = 68). **C** The fraction of females and males per plant for both sucrose treatments. Nematode sex ratios overturn between high sucrose and low sucrose conditions. Sex was scored 35 days after inoculation. Statistics are based on t-tests.

Next, we set out to quantify male development in *G. pallida*. In four independent experiments, the high sucrose concentration in the culture media led to a significant increase in the number of females, from an average of 1.0 female per plant on low-sucrose medium to 41.8 females per plant on high-sucrose medium (**Figure 1B**; t-test, p = 9*10^-14^). Conversely, male numbers dropped from 8.7 to 2.9 males per plant (**Figure 1B**; t-test, p = 1*10^-7^). This threefold decrease of male formation under high-sucrose conditions proves that the sex determination is impacted by the sucrose availability of the host. Moreover, since females constitute 94% of the adult population on high-sucrose medium and males 89% on low-sucrose medium (**Figure 1C**), these treatments effectively induce female and male development, respectively. We therefore used sucrose levels in the culture media to study the mechanisms underlying environmental sex determination (ESD) of *G. pallida*.

### Female development associates with extensive transcriptional and post-transcriptional regulations

To investigate the transcriptional response leading up to ESD in *G. pallida*, we conducted *in vitro* infection assays on low- and high-sucrose media. Since we were able to phenotypically identify the first female juveniles from 9 dpi onwards (**Figure 1A**), we inferred that sex determination occurred within the first 9 days of infection. Therefore, for both treatments infection sites were harvested from root tissue at 1, 3, 6, and 9 dpi. The experiment was quadruplicated, and the collected samples were used for transcriptome sequencing. After library preparations, 34 of the 36 samples passed quality control and were used for further analyses. Principal component analysis based on nematode gene expression, showed that the first principal component (PC1) captured 24.2% of the variance and PC2 captured 18.6% of the variance (**Figure 2a**). Both PC1 and PC2 indicated that the transcriptome changed over time. Another 10.8% of the variance was captured on PC3, which separated the pre-parasitic samples from the parasitic samples (**Figure 2a**). Notably, PC3 also seemed to reveal some of the variance induced by the sucrose treatments, as samples of the same treatment tended to group together at later time points of infection (6 and 9 dpi). This sucrose-induced signal in the data indicates that differentially expressed genes could be linked to sex determination in *G. pallida*.

**Figure 2.**
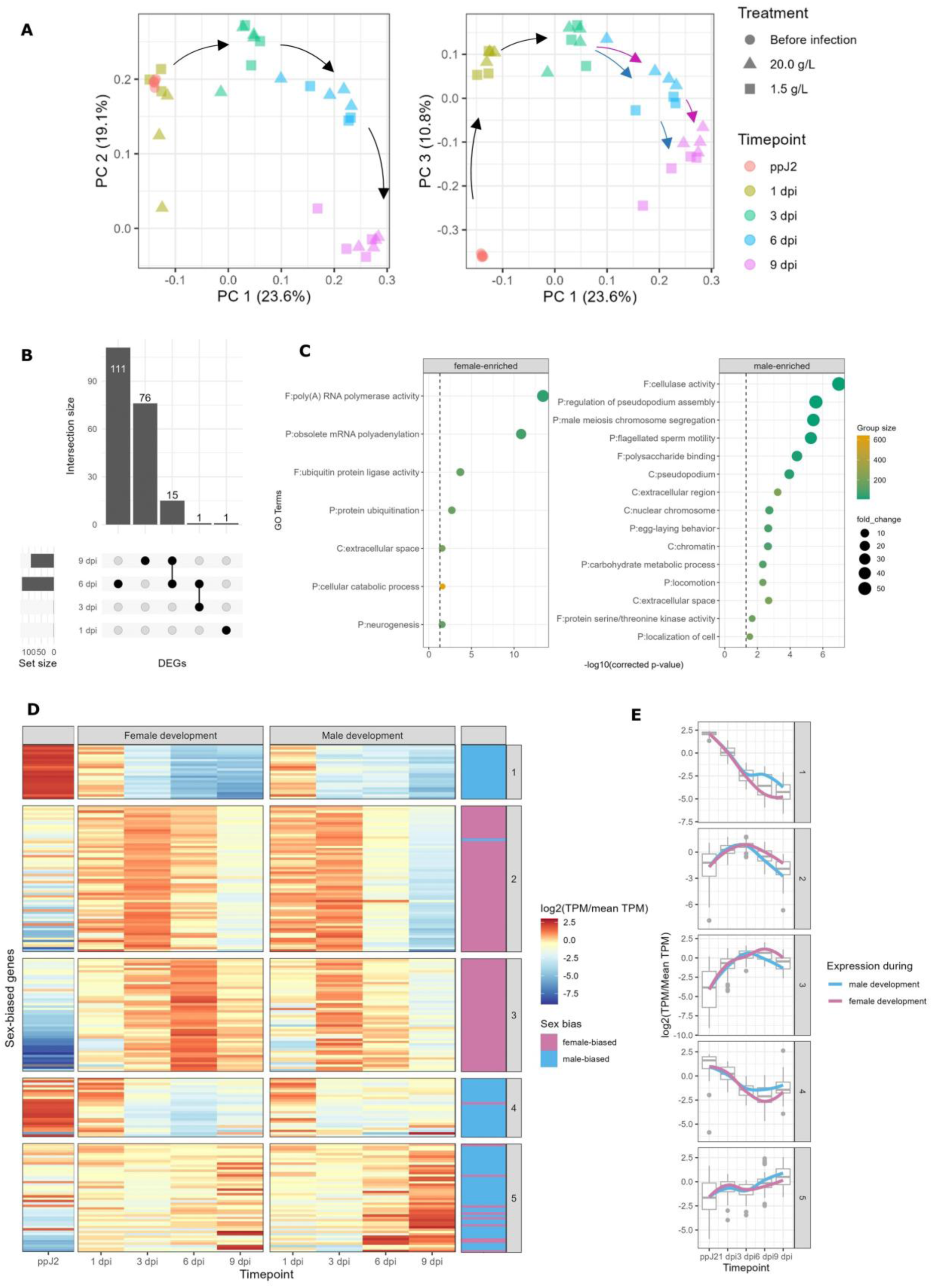
Transcriptomic differences indicate sex-biased clusters of differentially expressed genes. **A** Principal component analysis on the nematode gene expression. Principal component 1 (PC1) and PC2 associate with time. PC3 separates the pre-parasitic juveniles (ppJ2) from the parasitic juveniles. Moreover, from 6 dpi onwards samples cluster according to their sucrose treatment along PC3. Black arrows indicate the developmental direction over time. Pink and blue arrows indicate female- and male-biased development, respectively. **B** An UpSet plot indicating the distribution of the 204 differentially expressed genes (DEGs; FDR < 0.05) over time. Most DEGs were found at 6 and 9 dpi, as expected from the principal component analysis. **C** GO-term enrichment on female-biased and male-biased DEGs. The vertical line represents the significance threshold (FDR-corrected p-value = 0.05; -log_10_(p) = 1.3). Enrichment analysis was conducted on cellular components (C), biological processes (P), and Molecular functions (F). **D** A heatmap showing the relative expression of sex-biased genes during female and male development. K-means clustering clustered all DEGs into 5 distinct clusters based on their transcription over time. Expression is visualised as the log_2_ of the ratio between the TPM and the mean TPM. The column on the right indicates whether DEGs showed female- or male-biased expression. **E** Expression of DEGs over time per cluster. Line colours indicate sex-bias.

To identify nematode genes involved in ESD, we conducted differential expression analysis comparing high- and low-sucrose treatments at each timepoint. Out of the 18,606 genes, 204 unique genes (1.1%) were differentially expressed (FDR<0.05) between sucrose-treatments at one or more timepoints (**Figure 2b, Supplementary table 4**). Of these differentially expressed genes (DEGs), 85 genes were higher expressed in males, and 119 genes were higher expressed in females. Interestingly, the vast majority of all DEGs were identified at 6 and 9 dpi. At 1 and 3 dpi, only one gene showed differential expression between low- and high-sucrose treatments. This suggests that the development of males and females begins to diverge significantly between 3 and 6 dpi.

To assess whether the sucrose-induced DEGs were related to sex determination and sex differentiation, we analysed the gene ontology (GO) terms associated with the DEGs (**Supplementary table 4**). Of the 204 DEGs, 103 were annotated with GO terms. Among the annotated DEGs, 8 genes could be specifically linked to sex differentiation. Two genes involved in vulval development (*Gp_g6811*) and vitellogenesis (*Gp_6903*) were significantly higher expressed during female development. Conversely, three genes involved in male meiosis and sperm production (*Gp_g1313, Gp_g4334* & *Gp_g5290*), and three genes involved in muscle organ development (*Gp_g1284*, *Gp_g9515* & *Gp_g17790*) were significantly higher expressed during the development of males. GO-term enrichment analysis confirmed that male-biased genes were enriched for GO-terms related to male meiosis, sperm motility, and locomotion (**Figure 2C**). This includes two GO-terms related to pseudopodia, which are crucial for nematode sperm motility (Smith, 2014). However, the most significantly enriched GO-term among the male-biased DEGs was cellulase activity. This is in line with previous research that showed that cellulases were expressed in *Globodera* juveniles and males but not in females (Smant et al., 1997). Among the female-biased DEGs, genes associated with mRNA polyadenylation and protein ubiquitination were most significantly enriched (**Figure 2C**), suggesting that post-transcriptional regulation plays an important role in female development.

We observed that male-biased genes were enriched for genes involved in male sex differentiation and female-biased genes were enriched for genes related to post-transcriptional regulation. Therefore, we hypothesised that juveniles were predisposed to develop into males and only transition towards female development under favourable environmental conditions. To assess this hypothesis, we characterised the expression patterns of sex-biased genes over time. We conducted k-means clustering on the relative expression levels of all DEGs over time (**Figure 2D**). This separated all sex-biased genes into five clusters. The majority of male-biased genes cluster together in clusters 1 and 4, and the majority of female-biased genes cluster together in cluster 2 and 3. This indicates that male-biased genes and female-biased genes follow different transcriptional patterns over time. Male-biased genes were expressed from the pre-parasitic stage onwards and generally decrease in expression over time (**Figure 2E**). Conversely, female-biased genes were hardly expressed in the pre-parasitic stage and were mainly expressed once female development was initiated (**Figure 2E**). This supports the hypothesis of a default developmental trajectory towards males.

### Transcriptome profiling identifies three transcription factors as putative master regulators of environmental sex determination

Since transcription factors regulate sex in nematodes, mammals, and insects (Burtis & Baker, 1989; Gubbay et al., 1990; Hodgkin, 1987; Sinclair et al., 1990), we hypothesised that transcription factors also regulate ESD in *G. pallida*. Based on the GO terms, we identified three putative transcription factors among our DEGs (**Supplementary Figure 3A**), one of which (*Gp_g865*) shows female-biased expression (**Figure 3A**; FDR = 0.032) and is the closest related homolog of the zinc-finger transcription factor *lin-29* from *Caenorhabditis elegans* (BLASTp; 76% identity), a key regulator of developmental timing (Rougvie & Ambros, 1995). *Gp-lin-29* shares structural features with *Ce-lin-29,* including multiple c2h2 zinc finger domains, classifying it as part of the GLI family of zinc-finger proteins (**Supplementary Figure 3B**). Interestingly, in *C. elegans*, the expression of *lin-29* is also sex-biased, with significantly higher expression in males at the L1, L4, and young adult stages (Haque et al., 2024). Notably, *Gp-lin-29* also shares homology with *Ce-tra-1* (BLASTp; 43% identity), a key regulator of sex determination in *C. elegans* (Hodgkin, 1987). These similarities make *Gp-lin-29* an interesting candidate for involvement in ESD.

**Figure 3.**
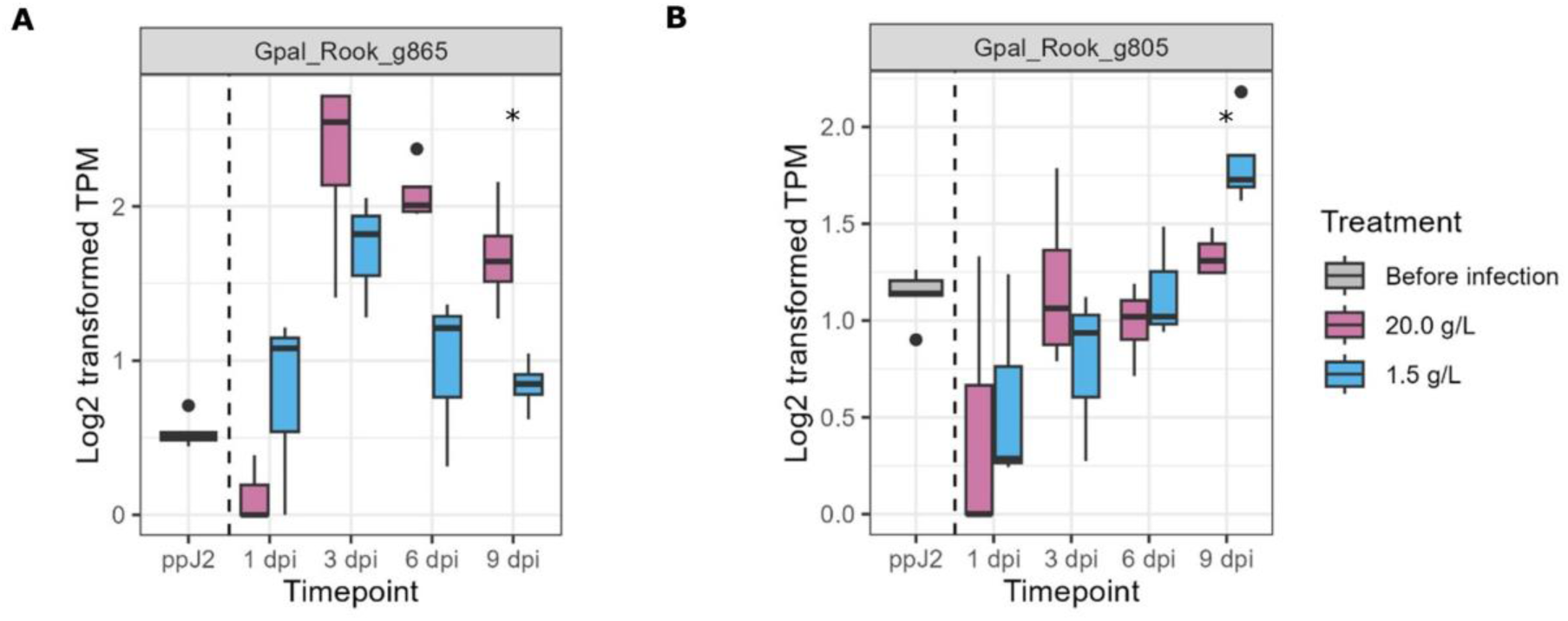
Differential expression of transcription factor *Gp-lin-29* and *Gp-mab-3*. Expression of genes associated with sex determination and sex differentiation during female (20 g/L) and male development (1.5 g/L). **A** Female-biased expression of *Gp-lin-29* at 9 dpi (FDR = 0.032), a putative transcription factor with 1-to-1 homology to *Ce-lin-29*. **B** The male-biased expression of *Gp-mab-3* at 9 dpi (p-value = 0.0045). *Gp-mab-3* is a putative regulator of sex differentiation with homology to *Ce-mab-3*. See **supplementary table 5** for more information on *G. pallida* homologs of components of the *C. elegans* somatic sex determination pathway.

Next, we assessed via which molecular pathways *Gp-lin-29* may be involved in the sex determination of *G. pallida*. Given *Gp-lin-29* showed homology with *tra-1* and displayed sex-biased expression, we hypothesized that components of the genetic sex determination pathway of *C. elegans* were present in nematode species exhibiting ESD. To test this, we screened cyst nematode genomes for homologs of seventeen known genes involved in the global somatic sex determination pathway in *C. elegans* (Haag, 2005). We identified homologs of five genes in *Globodera* (*fem-1*, *fem-2*, *fog-3*, *sel-10,* and *tra-3*; **Supplementary table 5**). However, none of these homologs showed differential expression between male and female development in *G. pallida* (**Supplementary Figure 4**). In addition, we looked for homologs of two genes regulated by *tra-1* that are involved in sex differentiation (**Supplementary table 5**). This includes the conserved transcription factor *mab-3*, for which the *G. pallida* homolog showed male-biased expression in *G. pallida* at 9 dpi (p-value = 0.0045, **Figure 3B**). As *mab-3* is critical for proper male formation in free-living and plant parasitic nematodes (Yi & Zarkower, 1999; Zhou et al., 2021), *Gp-mab-3* may be involved in sexual differentiation in *G. pallida* males. Taken together, this suggests that the *C. elegans* genetic sex determination pathway is not likely to be involved in ESD, but sexual differentiation pathways may be conserved.

In *C. elegans*, *lin-29* is known to be regulated by the conserved *let-7* miRNA (Pasquinelli et al., 2000; Rausch et al., 2015). Therefore, we first investigated whether we could identify the *Gp-let-7* miRNA within the *G. pallida* genome. Indeed, we found that *G. pallida* harbours a *let-7* homolog (*Gp_g6187*) that shares 100% identity with the mature *Ce-let-7* sequence. Therefore, we set out to investigate the role of small RNAs in ESD of *G. pallida*.

### Conserved miRNAs *let-7* and *miR-100* show sex-biased expression

To assess the role of small non-coding RNAs in ESD, all previously described RNA samples were also subjected to small RNA (sRNA) sequencing. This generated an average of 25.3 million clean reads per sample. The most abundant nematode reads were 22 and 23 nucleotides long with a guanine at the 5’-end (**Figure 4A**). Due to short reads being prone to ambiguous mapping, we aimed to minimize noise from potato-derived sRNA reads. Therefore, we only used ppJ2 reads to identify sRNA loci on the nematode genome with ShortStack (Johnson et al., 2016). This analysis yielded 45,894 sRNA loci with an average length of 245 bp. Principal component analysis based on the expression levels of all sRNA loci revealed that the primary source of variance among samples (PC1; 9.9%) associated with the difference between pre-parasitic and parasitic juveniles (**Figure 4B**). PC2 captured 7.6% of the variance and associated with time. PC3 captured 5.6% of the variance and roughly separated the male development samples from the female development samples (**Figure 4B**), indicating a clear sucrose-treatment-specific sRNA signal among the samples.

**Figure 4.**
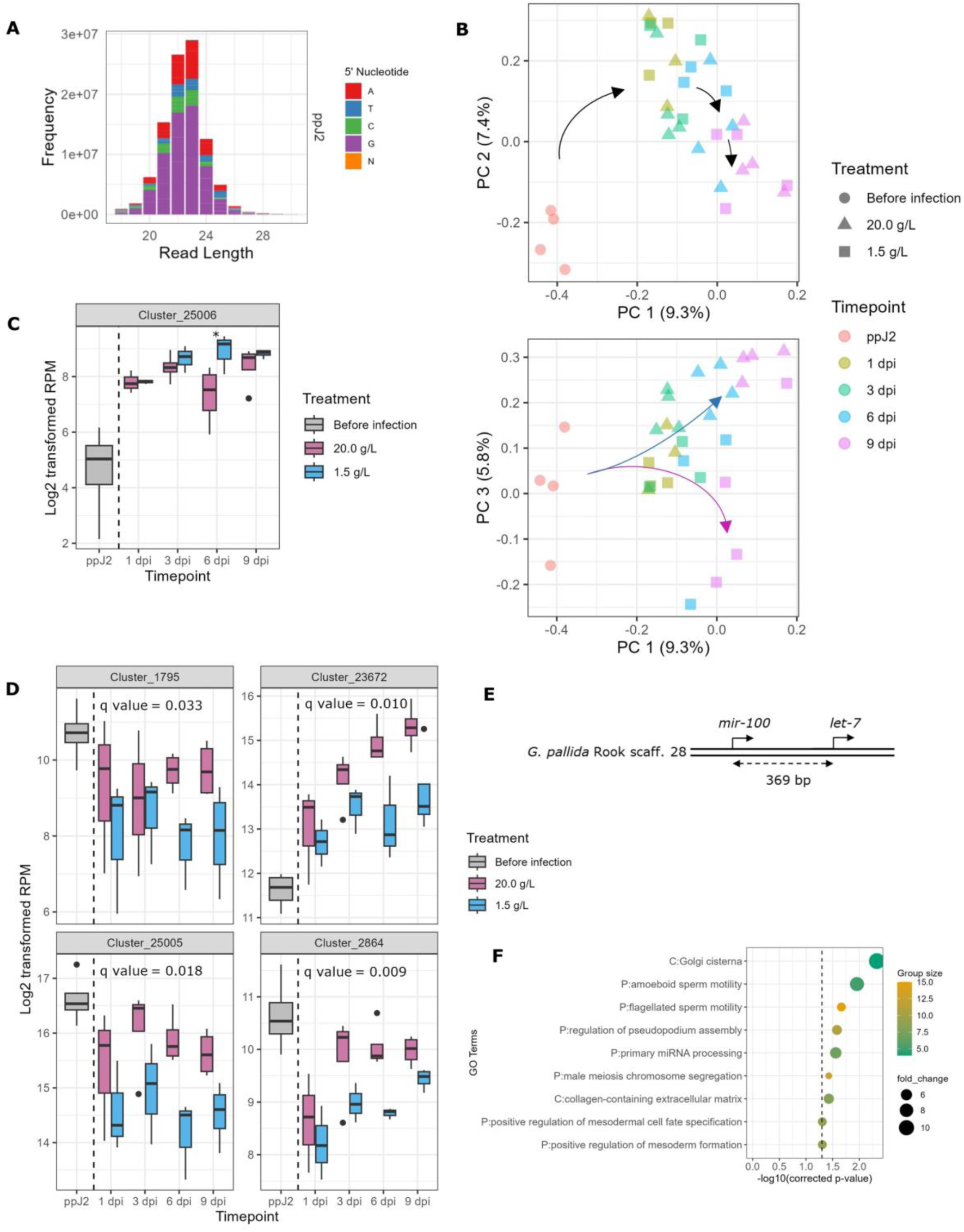
Conserved miRNAs are involved in environmental sex determination. **A** Principal component (PC) analysis on the nematode sRNA loci expression (RPM_ratio_). PC 1 splits pre-parasitic (ppJ2) samples and parasitic samples. PC 2 clusters samples by time. PC 3 clusters male samples and female samples together. **B** Read length distribution of sRNA reads mapping to the nematode genome. Colours indicate the 5’ nucleotide. Pre-parasitically, most reads are either 22 or 23G. **C** Expression of the miRNA *Gp-let-7*. Expression levels are significantly higher during male development (1.5 g/L sucrose) than during female development (20 g/L sucrose) at 6 dpi (p-value = 0.004). **D** The expression patterns of four miRNAs that are significantly higher expressed during female development than during male development (q-value < 0.05). The q-value is the FDR-corrected p-value. Cluster 25005 is the miRNA *miR-100*. Note that the y-axes do not start at zero. **E** Graphical representation of the *G. pallida let-7/miR-100* cluster. **F** GO-term enrichment analysis on the predicted miRNA targets of the four female-biased miRNAs identified nine significantly enriched GO-terms. MiRNA targets are enriched for cellular components (C), biological processes (P), and Molecular functions (F).

One of the predicted sRNA loci, cluster 25006 (Cl_25006), contained the *G. pallida let-7* homolog. Initially, this locus was not predicted as a miRNA locus by ShortStack. However, the secondary structure (**Supplementary Figure 5A**), a histogram of the read length distribution (**Supplementary Figure 5B**), a coverage plot of the locus (**Supplementary Figure 5C**), and the 100% identity with the *Ce-let-7* mature miRNA sequence strongly supports that this is the *Gp-let-7* miRNA. Interestingly, the *Gp-let-7* miRNA was up-regulated in males (p-value=0.005; **Figure 4C**). The subsequent down-regulation of *Gp-lin-29* (**Figure 3A**) supports our hypothesis that *let-7-*regulates expression of *lin-29*.

To evaluate the involvement of other sRNA loci in ESD, we compared all samples representing female development to all samples representing male development. Of the 45,894 loci identified, 326 (0.7%) were differentially expressed over time (FDR<0.05). Interestingly, 4 out of the 11 predicted miRNA loci (36%) showed differential expression, indicating that miRNA loci are significantly overrepresented among the differentially expressed loci (Hypergeometric test = 7.85e-07; **supplementary table 6**). All four miRNAs were higher expressed in females than in males (**Figure 4D**). Among these four miRNAs was the *miR-100* homolog (Cl_25005), located just 369 bp upstream of the *let-7* homolog (**Figure 4E**). *miR-100* is a member of the *let-7* cluster, which is conserved across many parasitic nematodes, flies, as well as humans (Pasquinelli et al., 2000; Sharma et al., 2024). The sex-biased expression of *miR-100* further supports a role for the involvement of *let-7* in the ESD mechanism.

### miRNA target prediction suggests that female-biased miRNAs suppress male developmental genes

To further investigate the role of the four female-biased miRNAs (Cl_25005, Cl_1795, Cl_2864 & Cl_23672), we aimed at identifying their target genes. Given that miRNAs primarily target mRNAs in the 3’ UTRs, we used the 500 bp downstream of each stop codon for miRNA target prediction (Majoros & Ohler, 2007). Based on the mature miRNA sequences, the RNAhybrid algorithm (Krüger & Rehmsmeier, 2006) identified between 9 and 212 putative targets with a minimal free energy below -25 kcal/mol and perfect seed matching per miRNA. Among the miRNA targets there were seven sex-biased DEGs (**Supplementary table 4**). The hits included male-biased genes associated with male meiosis, sperm development and multi-organism reproductive processes.

To further characterize possible functions of the predicted miRNA targets, we performed GO-term enrichment analyses. First, we conducted a GO-term enrichment analysis on the 610 predicted targets of all four female-biased miRNAs together. Nine GO-terms were found significantly enriched (**Figure 4F**). Four of these terms were related to sperm development, three of which were also found enriched among the male-biased genes (**Figure 2C**), suggesting that the primary role of these miRNAs was to suppress male development. The enrichment of ‘primary miRNA processing’ might indicate possible feedback loops for miRNA regulation. Second, we conducted separate GO-term enrichment analyses for the putative targets of individual miRNAs. In total, 20 unique GO-terms were found enriched (**Supplementary Figure 6**). The top GO-term for Cl_1795, ‘multi-organism reproductive process’, indicated that the putative targets of Cl_1795 were involved in processes related to sexual reproduction. Both Cl_1795 and Cl_25005 were enriched for GO-terms indicating regulatory roles at the genomic and transcriptional levels. The predicted targets of Cl_23672 were enriched for four splicing-related GO-terms as well as GO-terms related to the regulation of embryonic development. This connects to Cl_2864, which was enriched for three GO-terms associated with mesoderm formation and gastrulation. Although these processes primarily play a role in early embryonic development (Niu et al., 2022), these are also critical for proper body plan formation and gonadal tissue development (Ewen & Koopman, 2010). Together, this suggests that the four miRNAs are important for proper female development, primarily through suppression of male development.

## DISCUSSION

### Environmental sex determination provides an evolutionary advantage in *G. pallida*

Here, we demonstrate that favourable conditions (high-sucrose medium) promote female development, whereas unfavourable conditions (low-sucrose medium) promote male development in *G. pallida* juveniles. We show that an increased male-to-female ratio under nutrient-poor conditions is not merely attributable to stagnating female development (Von Sengbusch, 1927), but actually reflects a numeric increase in male formation. Therefore, we provide an even stronger indication for environmentally regulated sex. Our finding contributes to resolving a century-old debate surrounding sex determination in cyst nematodes. The first observation of alternating sex ratios in cyst nematodes was made in the year 1920 (Müller & Molz). The male-to-female ratio increased on nutrient-poor conditions, suggesting that sex cannot solely be a genetically predetermined trait. When the experiments were repeated, it was argued that the variable sex ratios resulted from an increased death rate of female nematodes in nutrient-poor conditions (Von Sengbusch, 1927). It was estimated that female nematodes need 35 times more nutrients to develop then male nematodes. Consequently, unfavourable conditions hamper female development and cause death. Although numerous observations were conducted on variable sex ratios in cyst nematodes ever since, the debate around the mechanism of sex determination in cyst nematodes remained unsettled. By measuring a numeric increase in males under nutrient-poor conditions we show that the host-derived nutrient supply is an important factor in the sex determination of cyst nematode juveniles (Anjam et al., 2020; Grundler et al., 1991). However, the exact nutritional cues that determine sex in *G. pallida* remain unknown.

We theorize that environmental sex determination (ESD) allows *G. pallida* to maximise its reproductive potential under favourable conditions, while maintaining its genetic variation under suboptimal conditions. If *G. pallida* was limited to GSD, many female juveniles would fail to reach maturity under unfavourable conditions and would be unable to contribute to the next generation. Whereas under favourable conditions developing as a female would maximize reproductive output, as females can outnumber the males because each male can mate with multiple females (Montarry et al., 2015). Thereby, ESD might provide *G. pallida* with a selective advantage over species constrained to GSD.

### *Globodera pallida* juveniles are predisposed to develop into males

The ability of cyst nematode juveniles to respond to external stimuli has been repeatedly discussed (Anjam et al., 2020; Grundler et al., 1991; Trudgill, 1967). Second-stage juveniles (J2s) were considered sexually undifferentiated until environmental cues direct their development towards a specific sex. However, our data indicates that J2s likely follow a default developmental trajectory leading to male development. Here we lay out the three arguments to support this hypothesis.

First, our transcriptomic analysis revealed that female-biased genes are generally up-regulated following infection, whereas male-biased genes are generally down-regulated over time. This suggests that female development requires transcriptional activation, while male development may proceed without extensive genetic reprogramming. Second, GO-term enrichment analysis on the sex-biased genes, indicated that female-biased genes are significantly enriched for post-transcriptional regulatory processes, including poly-A-modification and protein ubiquitination. Thus, female development requires extensive reprogramming through post-transcriptional modifications. Conversely, male-biased genes are not enriched for GO-terms related to gene regulation or post-transcriptional regulation. Third, all four miRNAs that showed to be sex-biased over the first nine days of infection are more abundant in females than in males, supporting the idea of a greater need for post-transcriptional regulation in female development. In addition, GO-term enrichment of the putative miRNA targets suggested that these female-biased miRNAs down-regulate genes involved in male development. This indicates that active suppression of male-specific genes is necessary for proper female development. Altogether, this makes it likely the default sex is male.

In addition, we theorize that a default male developmental trajectory is in line with resource availability. Namely, males arise in poor conditions, reducing the need for extensive reprogramming, making it a more efficient strategy. Alternatively, if juveniles had no default trajectory, all individuals would require transcriptional reprogramming to develop into either one of the sexes. Furthermore, if the default developmental program resulted in females, the resource-poor strategy (male development) would require an extra reprogramming effort. In conclusion, a male default would provide a resource-efficient path to maturation and reproductive contribution under nutrient-poor conditions.

### Identifying regulators of ESD is challenging, yet implicates a *lin-29*-like transcription factor

We identified sex-biased transcription factors by RNA sequencing a time series of infected root samples. Among the identified sex-biased transcription factors was *Gp-lin-29*, which shares homology with *Ce-tra1*, the master regulator of sex in *C. elegans* and insects (Beye et al., 2003; Chen et al., 2014; Hediger et al., 2010; Verhulst et al., 2010). This raised the question if *Gp-lin-29* could serve as a primary sex determination signal in *G. pallida*, potentially acting as the master regulator of ESD. Many well-characterised sex determination pathways are controlled by a master regulatory gene, such as the *Sry* gene in mammals (Foster & Graves, 1994), *dmrt1* in many other vertebrates (Smith et al., 2009; Volff et al., 2007; Yoshimoto et al., 2008). These conserved genes have evolved as primary sex determination signals, suggesting that certain transcription factors are particularly suited to regulate sex determination. The sequence identity and the domain conservation between *Gp-lin-29* and *tra-1*, makes *Gp-lin-29* an interesting candidate master regulator.

However, the identification of regulators of sex determination has proven to be challenging. Master regulators are not necessarily conserved transcription factors. For instance, *Sxl* in *Drosophila* (Maine et al., 1985) and *sdY* in *Oncorhynchus mykiss* (Yano et al., 2012) are non-conserved genes and do not encode transcription factors. It is likely that many non-conserved sex-determining genes have not been discovered yet. In the case of the specialized cyst nematodes, their evolutionary history suggests an ancestor that possessed genetic sex differentiation including a sex-chromosome (Gonzalez de la Rosa et al., 2020). It is therefore possible that neofunctionalization played a role in the selection for ESD. As only half of the sex-biased genes were functionally annotated with GO-terms and 76 out of 204 lacked informative BLASTp hits. A total of 71 sex-biased genes (35%) had no functional information available. Given the scope of this study, we were unable to investigate previously uncharacterized genes. The limited functional knowledge of the sex-biased genes further complicates data interpretation.

Alternative to the single-master-regulator model, ESD in *G. pallida* may be polygenic (Kocher et al., 2024). In several fish and vertebrate species, ESD is controlled by the combined effects of many interacting loci forming a quantitative threshold (Anderson et al., 2012; Bradley et al., 2011; Liew et al., 2012; Parnell & Streelman, 2013; Ser et al., 2010; Vandeputte et al., 2007; Yusa, 2007). When this model holds true for *G. pallida*, *Gp-lin-29* could be one of many genes contributing to a broader regulatory network. Third, the identification of sex determination genes in plant-parasitic nematodes based on RNA sequencing has been challenging for four technical reasons. First, despite generating over 100 million reads per sample, on average, only 4.8% mapped to the *G. pallida* genome, reducing our ability to detect lowly expressed regulatory genes. If a master regulator only functions in a small subset of cells, its expression may well have been below our detection threshold. Our data may primarily capture secondary signals coordinating sex differentiation. Second, since the exact timing of sex determination remains unknown, we may have missed the primary signal in the transcriptome. To address this, samples should be collected with shorter time intervals for the first nine days after inoculation to ensure temporal coverage at a higher resolution. Third, the infection sites collected for sequencing did not represent perfectly synchronised stages of infections. Although plants were inoculated simultaneously, some juveniles infected the roots within hours, while others took several days to infect, resulting in biological variation. Taken together, we must be aware of the challenges and shortcomings to interpret the data.

### A sex-biased cluster of miRNAs supports a role for *Gp-lin-29* in the sex determination of *G. pallida*

To investigate whether *Gp-lin-29* contributes to sex determination in *G. pallida*, we examined the abundance of the miRNA *let-7*, which is known to regulate *lin-29* in *C. elegans* (Pasquinelli et al., 2000; Rausch et al., 2015). Small RNA sequencing analysis revealed the male-biased expression of *let-7* in *G. pallida* at 6 dpi, and miRNA target prediction revealed *lin-29* as a putative target, supporting the hypothesis of a conserved regulatory role for *Gp-let-7*. In *Drosophila*, *let-7* has been identified as the primary modulator of the sex-determination hierarchy (Fagegaltier et al., 2014). Although *let-7* is up-regulated in both sexes during metamorphosis (Li et al., 2024), *let-7* levels were significantly higher in male gonads compared to female gonads (Fagegaltier et al., 2014). Functional studies have shown that *let-7* is essential for male sexual identity, as loss of *let-7* expression in males leads to the expression of genes that are normally restricted to females (Fagegaltier et al., 2014).

Like *tra-1* in *C. elegans*, *let-7* and *lin-29* may play a role at the interface of sex determination and sexual differentiation. *tra-1* is the terminal regulator of the sex determination pathway and it directs tissue-specific sex differentiation (Haag, 2005). It is expressed throughout the body but its effects on target genes are highly cell-type specific. Also, in *C. elegans*, *let-7* and *lin-29* influence the sex differentiation pathway to ensure correct sex-specific development (Sun & Hobert, 2023). For example, in male brains, *let-7* regulates sexually dimorphic neuron differentiation by targeting *lin-41* through *lin-29a* (Pereira et al., 2019). Since *let-7* itself is known to be hormonally regulated (Caygill & Johnston, 2008; Fagegaltier et al., 2014; Rubio et al., 2012; Song et al., 2018), future research may focus on the role of endocrine signalling in ESD.

We found another sex-biased miRNA, *Gp-miR-100*, just 369 base pairs upstream of *Gp-let-7*. The *miR-100* miRNA is highly conserved and among the most abundant miRNAs in free-living and plant-parasitic nematodes (Kaur et al., 2017; Sharma et al., 2024; Winter et al., 2012), although it is notably absent in *C. elegans* (Roush & Slack, 2008). In some organisms, *let-7* is co-expressed with *miR-100* as both miRNAs are derived from a common polycistronic transcript (Sokol et al., 2008). Our data shows that *Gp-let-7* is male-biased, while *Gp-miR-100* is female-biased, suggesting that the transcription of both miRNAs is regulated independently. However, since the accumulation rate of mature *let-7* and *miR-100* depends on the miRNA-specific processing rates of Drosha and Dicer (Chawla et al., 2016), differences in processing efficiency could result in sex-biased accumulation, even if both miRNAs are derived from the same transcript. In the free-living nematode *Pristionchus pacificus*, *miR-100* is developmentally restricted. Knockout mutants of *let-7* and *miR-100* showed developmental defects and reduced brood size (Sharma et al., 2024). Beyond nematodes, *miR-100* and *let-7* have also been implicated in sex-biased expression in embryos of species with a skewed male-to-female ratio, such as cows, yellow catfish and crabs, (Jing et al., 2014; Luo et al., 2021; Salilew-Wondim et al., 2024). The sex-biased expression of these miRNAs across diverse taxa suggests that their role in sex determination may be more widespread than previously recognized. We independently associated *let-7* and *miR-100* with sex determination, providing strong support for a functional role for this miRNA cluster in regulating ESD in *G. pallida*.

### The conserved *mab-3* and four sex-biased miRNAs associate with sex differentiation

Sex determination genes have been identified in many animals, but how the expression of these genes results in sexual dimorphism is poorly understood. We found male-biased expression of *Gp-mab-3*, a putative regulator of sex differentiation. In *C. elegans* the gene *mab-3* is one of the downstream targets of *tra-1* (Yi et al., 2000). The *mab-3* gene is also a conserved transcription factor (Panara et al., 2019) critical for proper male formation in free-living and plant parasitic nematodes (Yi & Zarkower, 1999; Zhou et al., 2021). In the pinewood nematode *Bursaphelenchus xylophilus*, *mab-3* was crucial for spermatogenesis, ontogenesis and mating behaviour (Zhou et al., 2021). Because of its high conservation, it is plausible that *mab-3* is also crucial for sexual differentiation in *G. pallida* males.

The identification of four female-biased miRNAs suggested that repressing male-specific genes is crucial for proper female development. When analysing the putative targets of these miRNAs, four sperm-related GO-terms were significantly enriched. Three of these sperm-related GO-terms were also enriched among the male-biased genes, further supporting a role for these miRNAs in the suppression of male-specific genes in female juveniles. This is comparable to how the *mir-35* family suppresses male-specific genes in *C. elegans* hermaphrodite embryos (McJunkin & Ambros, 2017). Further analysis of individual miRNA targets suggests regulatory roles in sexual reproduction (Cl_1795), body plan formation (Cl_2864), and splicing (Cl_23672), reinforcing our idea that miRNAs are key factors in sexual differentiation of *G. pallida*. Given that sex-specific alternative splicing critically contributes to sex determination in species such as *D. melanogaster* (Burtis & Baker, 1989; Kopp, 2012), the silkworm *Bombyx mori* (Kiuchi et al., 2014), and *Daphnia magna* (Kato et al., 2024), the miRNA Cl_23672 may similarly modulate sex-specific splicing in *G. pallida*. Future research employing long-read RNA sequencing could elucidate the role of alternative splicing in the ESD of cyst nematodes.

### Conserved components of the genetic sex determination pathway in *C. elegans* are absent in cyst nematodes

Phylogenetic analyses have not offered an explanation of the evolutionary origin of ESD in nematodes (Haag, 2005) and vertebrates (Navara, 2018). Frequent evolutionary transitions between GSD and ESD have led to the hypothesis that ESD develops when components in the GSD pathway become sensitive to environmental factors (Navara, 2018). This suggests that genetic sex determination pathways might be present in nematode species exhibiting ESD. A widely conserved general sex determination pathway is supported by the observation that some sex determination genes, such as *tra* and *her*, are conserved across phyla (Bachtrog et al., 2014). We found that the five (out of 17) *G. pallida* homologs we could identify (*fem-1*, *fem-2*, *fog-3*, *sel-10,* and *tra-*3) lack differential expression. This suggests that the genetic sex determination pathway as described in *C. elegans* is not likely to be involved in ESD. Rather, we hypothesize that the environmentally-regulated sex determination pathway is independent of the GSD pathway of *C. elegans*.

## Conclusions

Our findings provide evidence that the sex of *G. pallida* is environmentally regulated and strongly supports the hypothesis of a default developmental trajectory towards male differentiation. Upon establishment of a feeding site, nutritional cues can activate a signalling cascade that overrides this default pathway, enabling female development. This process requires extensive transcriptional activation and post-transcriptional regulation. A process possibly mediated by the transcription factor *Gp-lin-29*. The male-biased expression of its presumed suppressor *Gp-let-7* supports a role for this transcription factor in ESD. Additionally, female development requires the suppression of male-specific genes, a process facilitated by miRNAs that target male developmental genes. Future studies can use the high- and low-sucrose treatments to unravel the precise molecular mechanisms underlying sex determination in *G. pallida* and other cyst nematodes. Understanding these mechanisms may not only provide insight into nematode-host interactions, but also shed light on fundamental principles of developmental plasticity in larger animals.

## Author contributions

ASS, GS and MGS designed the experiments. ASS conducted the lab experiments. SJSvdR and ASS processed the sequencing data. ASS conducted data analyses. ASS, GS and MGS wrote the paper with input from SJSvdR.

## Acknowledgments

ASS thanks the EMBL-EBI (Hinxton, UK) for the course *‘Introduction to RNA-seq and functional interpretation’*, in which many of the software used in this study was introduced.

## Data availability

All scripts and underlying datasets are available through gitlab (https://git.wur.nl/published_papers/Schaveling_2025_ESDpallida). Raw sRNA sequencing data is deposited at BioStudies and available under accession code E-MTAB-15069, respectively. The sRNA loci annotation is deposited with the *G. pallida* Rookmaker genome at the NCBI (PRJEB91928).

## SUPPLEMENTARY FIGURES

**Supplementary figure 1.**
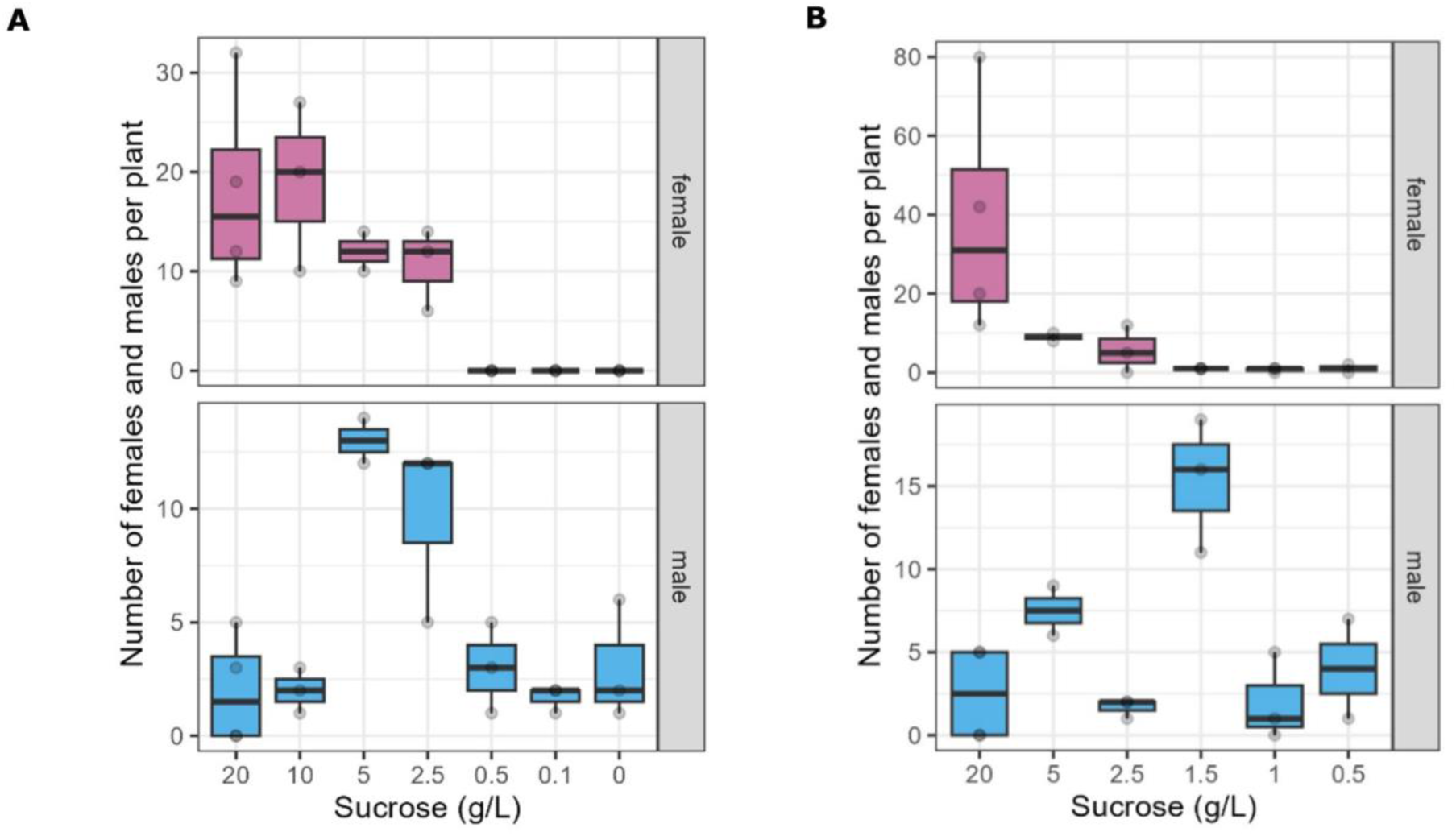
Data from two experiments to establish conditions in which the sex-determination of *G. pallida* can be manipulated. Potato cuttings were grown *in vitro* on Gamborg B5 medium supplemented with a range of sucrose concentrations. Males and females were scored at 35 dpi. **A** Trial 1 indicated that sucrose concentrations between 2.5 and 20 g/L allow for female formation, and sucrose concentrations above 0.5 and below 10 g/L promoted male formation. **B** Trial 2 indicated that 1.5 g/L sucrose induced the highest number of males, while hardly any females were formed at this concentration. Note that the scales are different between the panels and that both experiments were performed with a different batch of hatched juveniles.

**Supplementary figure 2.**
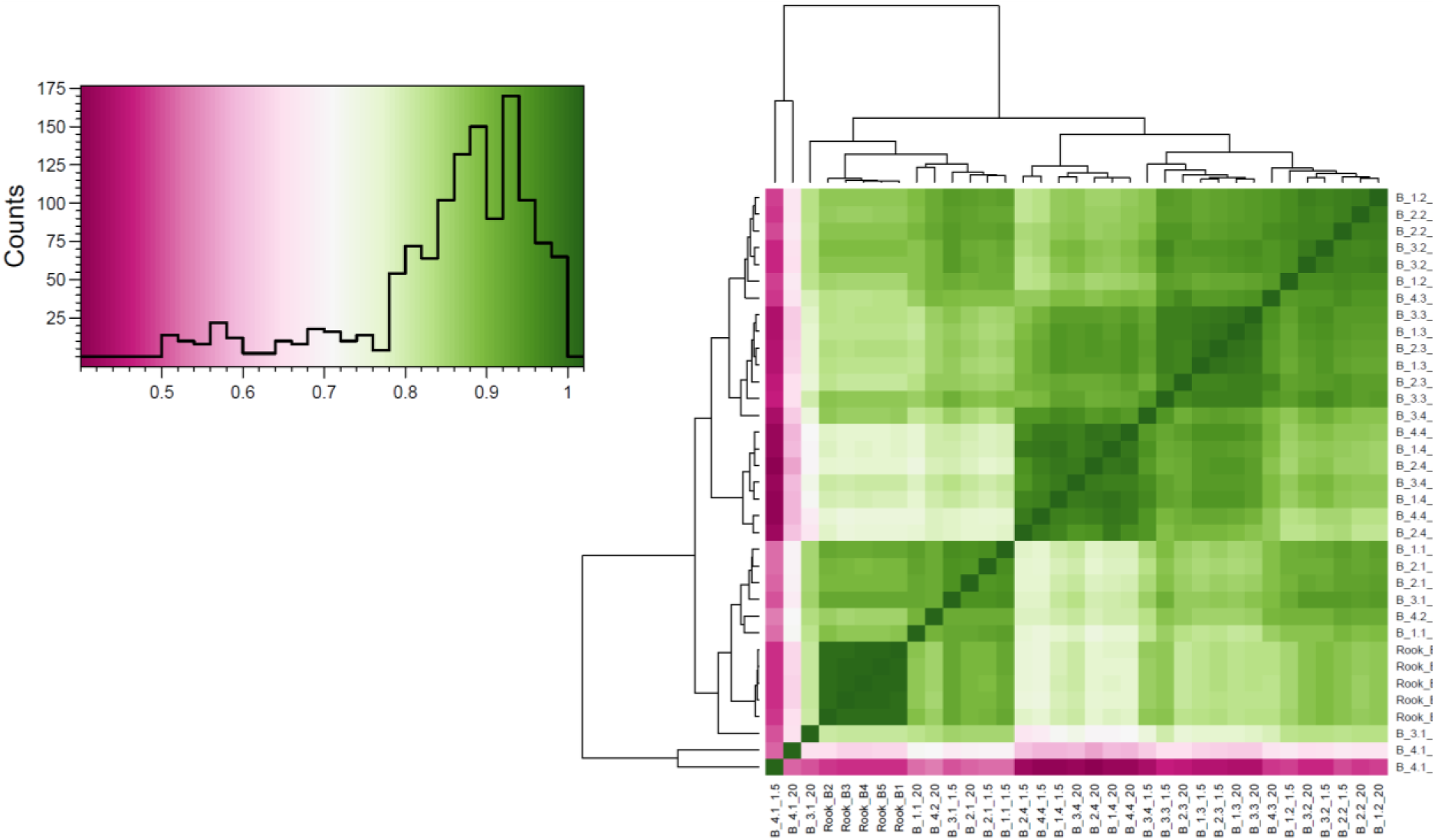
Pairwise Pearson correlations correlation matrix among all mRNA sequencing samples based on the log_2_ of the transcript per kilobase million (TPM) values. Based on the poor correlation both samples from batch 4 harvested at 1 dpi (B_4.1_20 and B_4.1_1.5) were excluded from further analyses.

**Supplementary figure 3.**
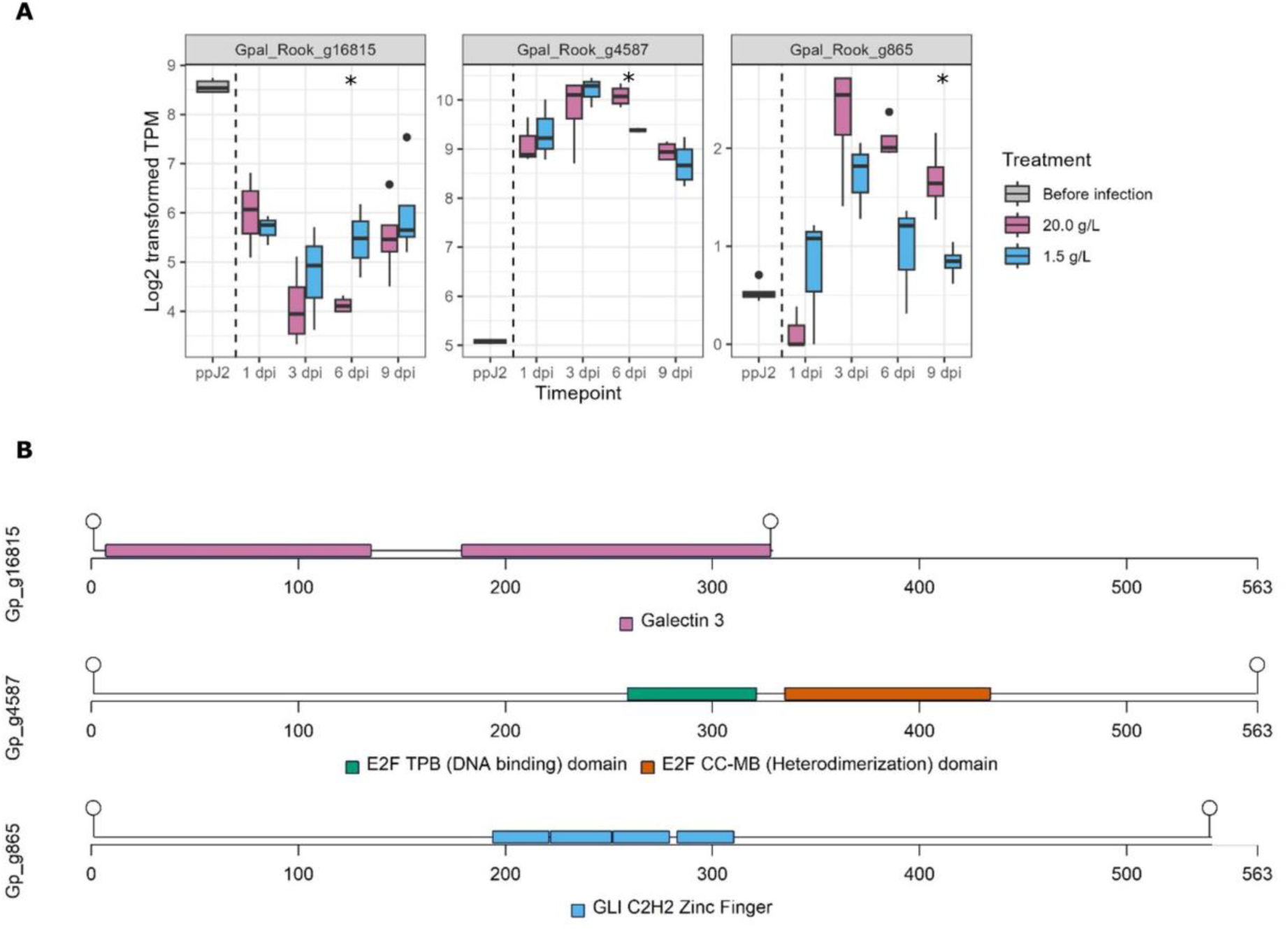
A. Expression of three putative transcription factors that were differentially expressed between male development (1.5 g/L sucrose) and female development (20 g/L sucrose). **B** A graphical representation of the three transcription factors showing their representative domains as predicted by InterPro (Paysan-Lafosse et al., 2023). Lollipops indicate protein start and end. The first transcription factor is Gpal_Rook_g16815 (*Gp_g16815*) and has two predicted sugar-binding pockets (Galactin-3 domains), suggesting a role in converting the metabolic signals into transcriptional regulation. However, since this transcription factor lacks a DNA binding domain, its function may depend on the dimerizing with other TFs. The second transcription factor is *Gp_g4587* and belongs to the E2 promotor-binding factor (E2F) transcription factor family. It features two distinct functional domains. The first is an E2F TDP domain, which facilitates dimerization with dimerization proteins (DP), forming an E2F-DP complex that regulates the cell cycle (Saad et al., 2021). The second is an E2F coiled-coil marked box (CC-MB) domain, which interacts with cyclin-dependent kinases (CDKs; Saad et al., 2021). The third TFs is *Gp-lin-29* and shares structural features with *Ce-lin-29,* including multiple c2h2 zinc finger domains, classifying it as part of the GLI family of zinc finger proteins.

**Supplementary figure 4.**
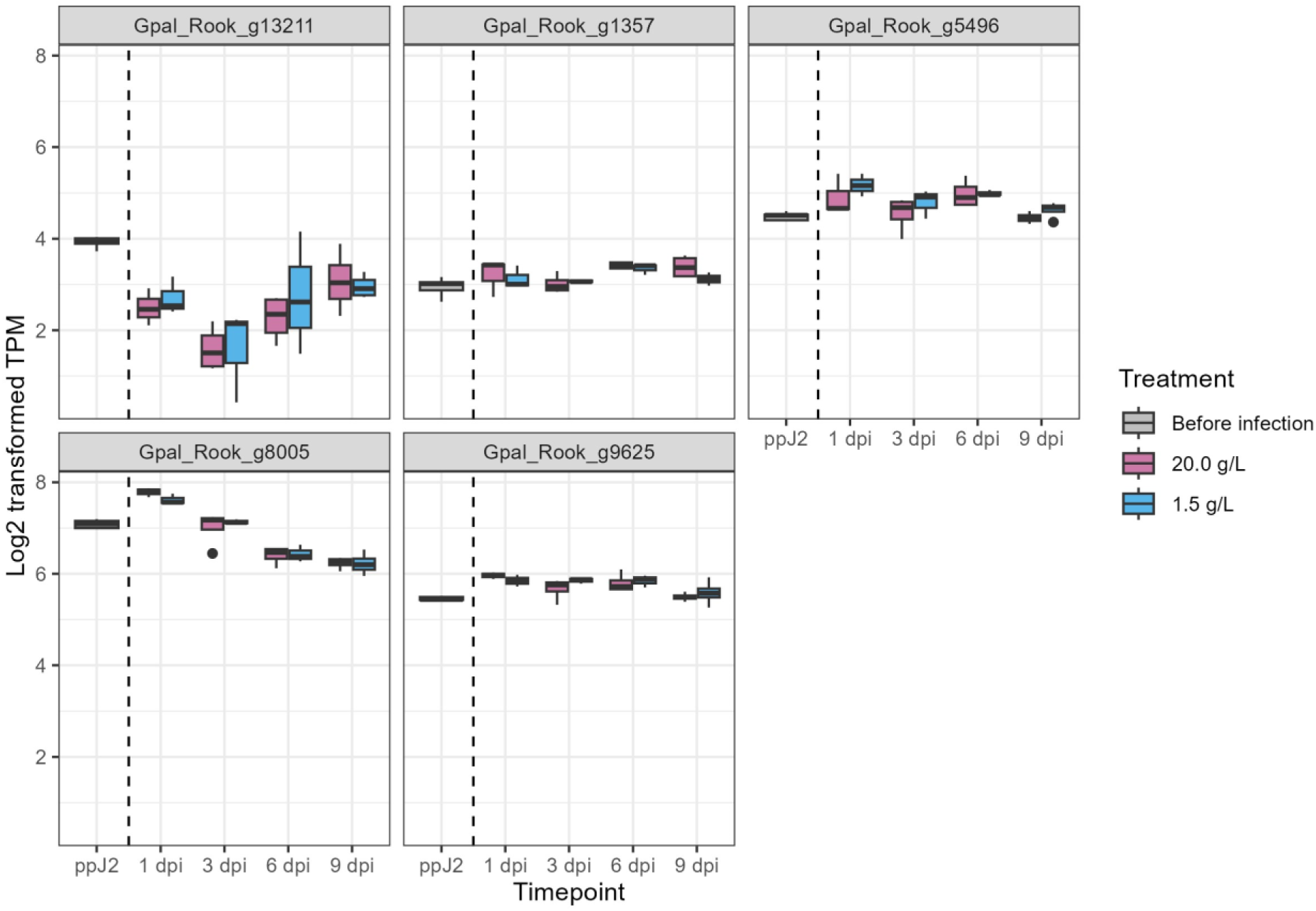
The expression of five *G. pallida* homologs of components of the somatic sex determination pathway of *C. elegans*: *tra-3* (Gpal_Rook_g13211), *fog-3* (Gpal_Rook_g1357), *fem-2* (Gpal_Rook_g5496), *fem-1* (Gpal_Rook_g8005), and *sel-10* (Gpal_Rook_g9625). See **supplementary table 5** for more information on *G. pallida* homologs of components of the *C. elegans* somatic sex determination pathway.

**Supplementary figure 5.**
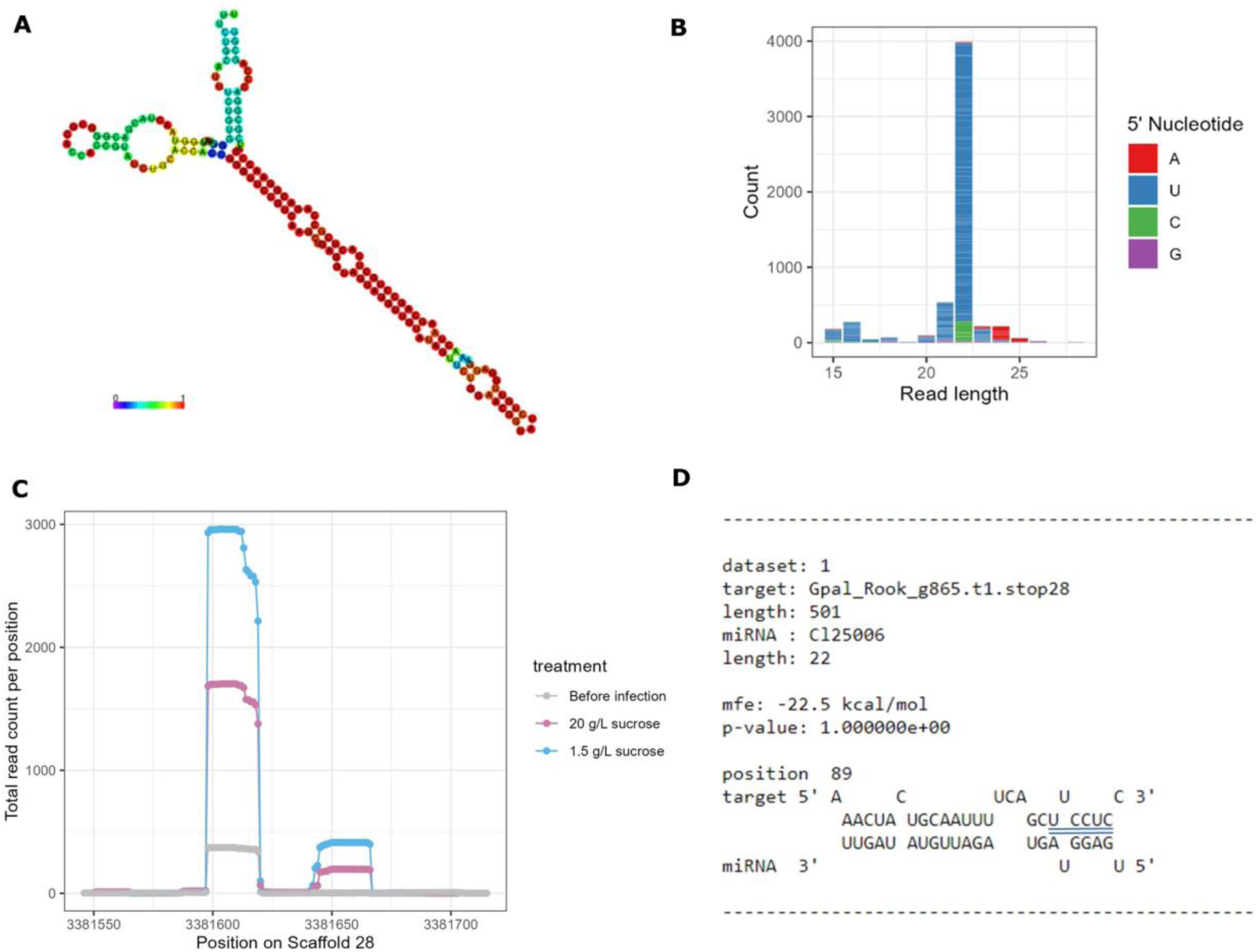
Characterisation of *Gp-let-7* **A** Predicted 2D structure of *Gp-let-7*. Colours indicate base-pair probabilities. **B** Read length distribution of small RNA reads mapping to *Gp-let-7*. Reads mapping to this locus mainly consist of the 22’U reads. **C** Coverage of the *Gp-let-7* locus, indicating the position of the mature miRNA sequence (left) and the miRNA star (right) **D** *Gp-lin-29* is predicted as a target of *Gp-let-7*, based on the minimal free energy (mfe) between the 3’UTR of *Gp-lin-29* and the mature miRNA sequence of *Gp-let-7*. It should be noted that there is one mismatch in the seed sequence. The seed sequence is highlighted by the blue lines.

**Supplementary figure 6.**
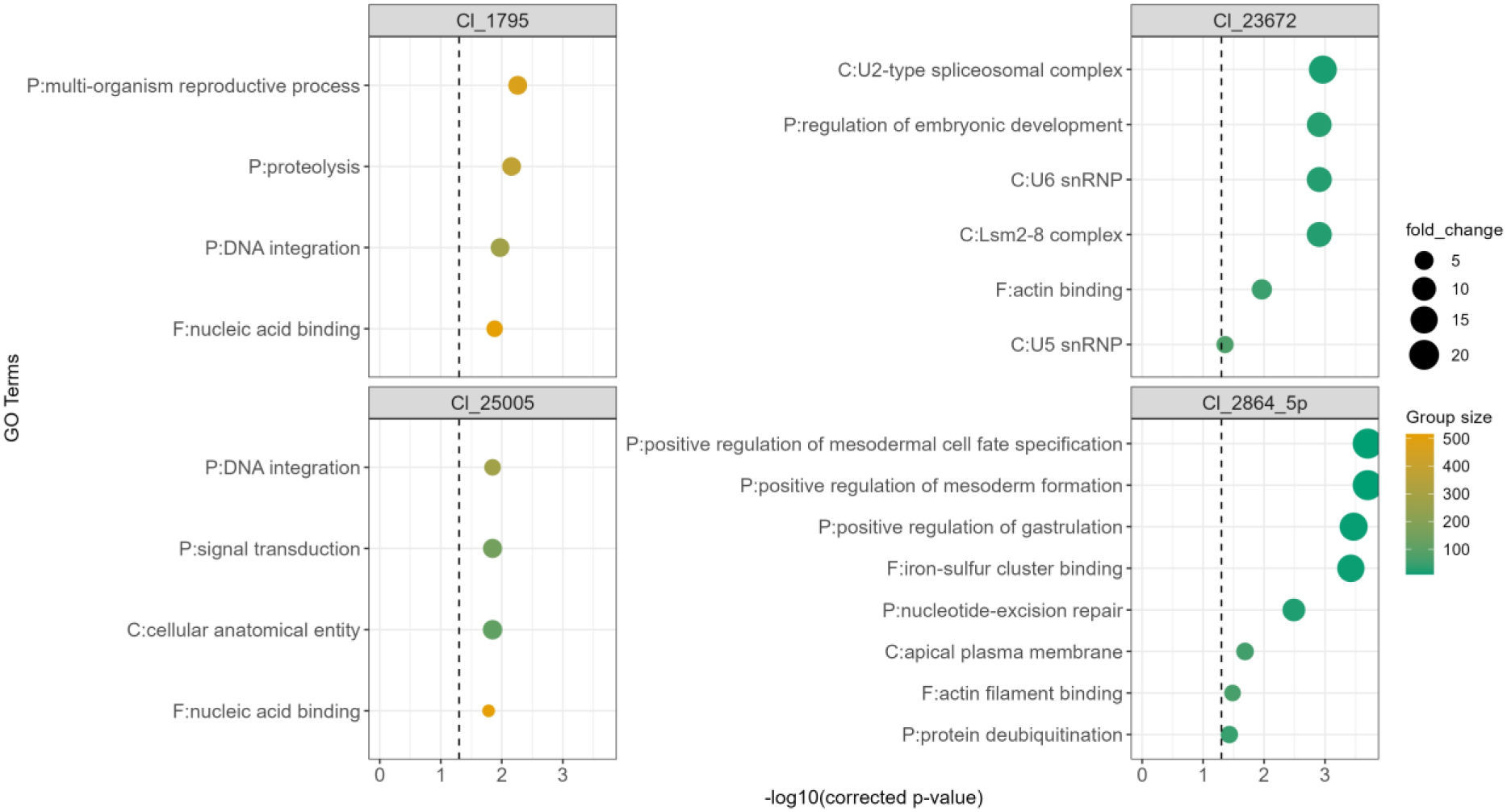
GO-term enrichment analysis on the predicted miRNA targets of each of the four female-biased miRNAs. Cluster 25005 is *Gp-miR-100*.

## SUPPLEMENTARY TABLES

An overview of all samples subjected to transcriptome sequencing (**Supplementary table 1**) and small RNA sequencing (**Supplementary table 2**). For each sample, read count and mapping statistics are provided. The two samples marked in red were not used in the analyses because of poor clustering (**Supplementary Figure 2**).

**Supplementary table 3**. An overview of for small RNA samples in which a more detailed output is given for the mapping statistics on the *G. pallida* Rookmaker genome.

**Supplementary table 4.** An overview of the 204 differentially expressed genes, including GO annotations, their best informative BLASTp hits. When DEGs are a predicted target of any of the sex-biased miRNAs, the miRNA is indicated.

**Supplementary table 5.** BLASTp hits of components of the *Caenorhabditis elegans* sex determination pathway on four cyst nematode genomes. BLASTp hits with an identity over 25% and a query coverage over 50% are considered homologs.

**Supplementary table 6.** An overview of the five sex-biased miRNAs found in this study, including their position, sequences, the number of putative targets and their closest homolog.

## SUPPLEMENTARY VIDEO

**Supplementary video 1.** A stop-motion video of a *G. pallida* juvenile developing into a female over the first 31 days post inoculation (dpi). At 31 dpi, a male is visible that is supposedly looking to mate with the female.

